# Excessive cholesterol accumulation in microglia increases neuronal synaptic vulnerability to amyloid-beta

**DOI:** 10.64898/2026.08.27.747668

**Authors:** Shihui Ding, Nicolaus Nazarenkov, Jungsu Kim, Kim Dore, Soo-Ho Choi, Yury I. Miller

## Abstract

Cholesterol efflux is an important determinant of cellular lipid homeostasis. However, how microglial excessive cholesterol accumulation affects neuronal synaptic integrity remains poorly understood, particularly in the context of Alzheimer’s disease. Here, we utilized a conditional knockout mouse model targeting the cholesterol transporters ABCA1 and ABCG1 in microglia. The microglia-specific ABCA1/ABCG1 deficiency triggered marked cholesterol accumulation, microglial hypertrophy, downregulation of the homeostatic marker *P2ry12*, and upregulation of the reactivity-associated marker CD11b, indicating shift toward a reactive phenotype. This phenotype was accompanied by increased reactive oxygen species, consistent with enhanced oxidative stress in ABCA1/ABCG1-deficient microglia compared with control. Using organotypic hippocampal slice cultures, we investigated the downstream neuronal outcomes of microglial ABCA1/ABCG1 deficiency. Under basal conditions, microglial ABCA1/ABCG1 knockdown did not significantly alter dendritic spine density in CA1 pyramidal neurons. However, upon exposure to amyloid-beta (Aβ) stress, microglial ABCA1/ABCG1 deficiency markedly exacerbated dendritic spine loss in CA1 pyramidal neurons. Taken together, our findings highlight an important role for ABCA1/ABCG1-dependent cholesterol efflux in maintaining microglial homeostasis and limiting neuronal synaptic vulnerability to Aβ-associated stress. These results support further investigation of microglial cholesterol transport as a potential target for preserving synaptic resilience in Alzheimer’s disease.

## Introduction

Cholesterol is an essential component of the brain and plays important roles in neuronal and synaptic function. Perturbation of cholesterol homeostasis can alter synaptic transmission and plasticity (1,2), and dysregulated cholesterol metabolism has been implicated in neurodegenerative disorders, including Alzheimer’s disease (AD), in which amyloid-beta (Aβ) accumulation is a characteristic pathological feature (3). ATP-binding cassette transporters ABCA1 (ATP-binding cassette subfamily A member 1) and ABCG1 (ATP-binding cassette subfamily G member 1) are major regulators of cellular cholesterol efflux (4,5) and act synergistically to facilitate cellular cholesterol removal (6). In the central nervous system (CNS), ABCA1 and ABCG1 participate in cholesterol handling across different cell types (7,8).

ABCA1 has also been linked to Aβ-related pathology in mouse models of AD. ABCA1 deficiency increased amyloid deposition in PDAPP and APP23 mice, although its effects on amyloid burden varied among AD models (9–11). Conversely, brain overexpression of *Abca1* reduced Aβ deposition and amyloid plaque burden in PDAPP mice (12). These studies suggest that ABCA1-dependent lipid homeostasis can influence Aβ-related pathology. Beyond its association with amyloid pathology, ABCA1 has also been implicated in neuronal and synaptic integrity. Brain-specific *Abca1* knockout mice exhibited reduced total and excitatory synapse density, together with decreased synaptic vesicle density (13). Moreover, brain-specific *Abca1* knockout mice developed increased cortical neuronal death at advanced age, whereas comparable neuronal death was not observed in mice following selective deletion of *Abca1* in neurons or astrocytes (14). ABCG1 has also been implicated in neuronal integrity, as CNS ABCG1 deficiency exacerbated neuronal injury and neurological deficits following traumatic brain injury (15). Together, these findings implicate ABCA1- and ABCG1-dependent cholesterol homeostasis in neuronal integrity, but the contribution of individual CNS cell populations remains unclear.

ABCA1/ABCG1-dependent cholesterol transport has also been shown to regulate microglial function. In a neuropathic pain model, ABCA1/ABCG1 deficiency in spinal microglia increased lipid raft abundance and TLR4 dimerization and induced basal allodynia, whereas cholesterol depletion with 2-hydroxypropyl-β-cyclodextrin alleviated this phenotype (16,17). These findings support a functional role of ABCA1/ABCG1-dependent cholesterol homeostasis in microglia.

Despite these observations, whether ABCA1/ABCG1-dependent cholesterol homeostasis in brain microglia contributes to neuronal synaptic integrity remains unclear. In particular, it is unknown whether impaired microglial cholesterol transport alters synaptic vulnerability to Aβ-associated stress. Here, we used an inducible microglial ABCA1/ABCG1-deficient mouse model to investigate how disruption of microglial cholesterol transport affects microglial state, and whether these changes influence dendritic spine integrity under basal and Aβ-stressed conditions. We found that microglial ABCA1/ABCG1 deficiency resulted in cholesterol accumulation, promoted a reactive microglial phenotype accompanied by increased oxidative stress, and enhanced neuronal susceptibility to Aβ-induced dendritic spine loss.

## Materials and Methods

### Animals

*Abca1^fl/fl^ Abcg1^fl/fl^ Cx3cr1-Cre^ERT2^* (ABC-imKD) mice were generated as previously described (17), and *Abca1^fl/fl^ Abcg1^fl/fl^* mice were used as WT controls. The *Cx3cr1-Cre^ERT2^* (Litt) line was obtained from The Jackson Laboratory (JAX Stock No. 021160), and subsequent offspring were used for experiments. All mice were on a C57BL/6J background. Mice were housed at room temperature in standard laboratory cages (up to five per standard cage) under a 12–h light/12–h dark cycle, with food and water provided ad libitum. All experiments included mice of both sexes and were performed in accordance with protocols approved by the Institutional Animal Care and Use Committee of the University of California, San Diego.

### Neonatal intragastric administration of tamoxifen for CreERT2 induction

A tamoxifen (TAM) stock solution (5 mg/ml in corn oil) was prepared by shaking overnight at 37 °C, protected from light, and stored at 4 °C for no longer than 2 weeks. Neonatal mice received 50 µg TAM per pup or vehicle (corn oil) by intragastric administration once daily on postnatal days 1–3 (P1–P3). A 1 mL syringe (BD, 309659) fitted with a 30-G × 1/2-inch needle (BD, 305106) was used for intragastric administration.

### Brain section preparation

Postnatal pups (P7–P13) were anesthetized by intraperitoneal injection of a mixture of ketamine (100 μg/g) and xylazine (10 μg/g), followed by transcardial perfusion with PBS and freshly prepared 4% paraformaldehyde (PFA). Brains were post-fixed overnight in 4% PFA at 4 °C and subsequently cryoprotected in a graded sucrose series (10%, 20% and 30% in PBS). After embedding in OCT compound, brain blocks were flash-frozen in liquid nitrogen and stored at −80 °C. Brain blocks were sectioned serially at 20 μm thickness using a cryostat, and sections were stored at −20 °C before use.

### Immunofluorescence labeling of brain sections

Brain sections were washed with Tris-buffered saline (TBS), permeabilized with 0.2% Triton X-100 in TBS (TBS-T), and blocked with blocking buffer containing 3% BSA and 3% normal goat serum in TBS-T. Sections were incubated with primary antibodies overnight at 4 °C. After three washes with TBS-T, sections were incubated with secondary antibodies for 1–2 h at room temperature. When applicable, DAPI was included after the secondary antibody incubation step for nuclear staining. Following additional washing steps, sections were treated with TrueBlack autofluorescence quencher (Biotium, 23014), washed several times with TBS, and mounted with ProLong Glass Antifade Mountant (Invitrogen, P36984). Primary antibodies used were anti-ABCA1 (Novus Biological, NB400-105, 1:200), anti-ABCG1 (Proteintech, 13578-1-AP, 1:200) and red fluorochrome (635)-conjugated anti-IBA1 (Fujifilm, 013-26471, 1:500). The secondary antibody used was goat anti-rabbit IgG conjugated to Alexa Fluor 568 (Invitrogen, A-11011, 1:1000). The fluorophore-conjugated anti-IBA1 antibody was applied after the secondary antibody incubation.

### Imaging acquisition and analysis

Fluorescence images of stained mouse brain sections were acquired using a Leica SP8 confocal microscope equipped with a ×20/0.75 NA dry objective (1024 × 1024 pixels). Image analysis was performed using Imaris software. Microglial cell boundaries were segmented using the Surface function based on IBA1 fluorescence. The area of IBA1^+^ cells was calculated based on the segmentation. The median cell area was calculated for each mouse and used as the biological replicate for statistical analysis. For each mouse, 55–118 IBA1^+^ microglia from the hippocampal region were analyzed for ABCA1 or ABCG1 mean fluorescence intensity (MFI), and the median MFI across all analyzed microglia was calculated. Background fluorescence was determined from brain sections processed in parallel without incubation with anti-ABCA1 or anti-ABCG1 primary antibodies. The batch-specific background median MFI was subtracted from the median microglial MFI to obtain the corrected median MFI for each mouse. All image processing and quantification were carried out in a blinded manner. Statistical analyses were conducted using GraphPad Prism version 10.

### Isolation of postnatal microglia-enriched brain cells

Microglia were isolated using a method modified from previous studies (18–20). Postnatal pups were anesthetized by intraperitoneal injection of ketamine/xylazine and decapitated. Whole brains were mechanically dissociated using a tissue grinder (EMS, 64793-72), and the resulting cell suspension was subjected to 37%/70% Percoll (Sigma-Aldrich, P4937) density gradient centrifugation to enrich for microglia.

### Analysis of ABCA1 expression by flow cytometry

For ABCA1 expression analysis, microglia-enriched cell suspensions were stained with Ghost dye red 780 (CST, 18452) to exclude dead cells and then fixed with freshly prepared 4% PFA for 10 min. Following fixation, cells were quenched with 1.5 mg/ml glycine in PBS. Cells were then blocked in PBS containing 2% anti-mouse CD16/CD32 (Mouse BD Fc Block, 553141, 1:50), 2% normal mouse serum, 1% BSA, and 0.1% Saponin. Cells were subsequently stained with anti-CD11b-PE/Cy7 antibody (Biolegend, 101216, 1:50), anti-ABCA1 antibody (Novus Biological, NB400-105, 1:50), and Alexa Fluor 568-conjugated anti-rabbit secondary antibody (Invitrogen, A-11011, 1:500). After washing with PBS containing 1% BSA and 0.1% Saponin, samples were analyzed on a CytoFLEX flow cytometer (Beckman Coulter), and data were analyzed using FlowJo software (BD Bioscience). ABCA1 fluorescence was quantified as the median fluorescence intensity in ABCA1^+^CD11b^+^ live singlets using the ECD channel.

### Filipin staining

Filipin III was used to detect cellular unesterified cholesterol. Filipin staining was performed using protocols modified from previous studies (21–23). For filipin staining of brain sections, frozen sections were washed with PBS, incubated with 1.5 mg/ml glycine in PBS for 10 min, and stained with 50 μg/ml filipin III (Sigma-Aldrich, F4767) for 2 h at room temperature in the dark. After three washes with PBS, sections were mounted with ProLong Glass Antifade Mountant (Invitrogen, P36984). Confocal images were acquired using the DAPI channel. Because the *Cx3cr1-Cre^ERT2^* line (JAX Stock No. 021160) express a Cre-ERT2 fusion protein and EYFP from the endogenous *Cx3cr1* promoter/enhancer elements, EYFP fluorescence was used to identify *Cx3cr1*-expressing microglia in postnatal brain sections (24).

For filipin staining of microglia-enriched cell suspensions, the suspensions were stained with Ghost dye red 780 (CST, 18452) to exclude dead cells, then fixed with freshly prepared 4% PFA for 10 min, and quenched with 1.5 mg/ml glycine in PBS. Cells were then blocked with PBS containing 2% anti-mouse CD16/CD32 (Mouse BD Fc Block, 553141, 1:50) and 2% normal mouse serum, labelled with anti-CD11b-PE/Cy7 antibody (Biolegend, 101216, 1:50), and stained with 100 μg/ml filipin (0.4% DMSO) for 1 h at room temperature in the dark. After two washes with PBS, samples were analyzed on a CytoFLEX flow cytometer (Beckman Coulter), and data were analyzed using FlowJo software (BD Bioscience). Filipin fluorescence was quantified in CD11b^+^ microglia using the PB450 channel.

### DHE staining

Dihydroethidium (DHE) fluorescence was used as an indicator of intracellular superoxide-associated oxidative stress. DHE staining was performed using a protocol adapted from a previous study (25). Microglia-enriched cell suspensions were blocked with PBS containing 2% anti-mouse CD16/CD32 (Mouse BD Fc Block, 553141, 1:50), labelled with anti-CD11b-APC antibody (BD, 553312, 1:50), and incubated with 5 μM DHE in HBSS for 10 min at 37 °C in the dark. After two washes with PBS, DAPI was added immediately before flow cytometric analysis to exclude nonviable cells. Samples were acquired on a CytoFLEX flow cytometer (Beckman Coulter), and data were analyzed using FlowJo software (BD Bioscience). DHE fluorescence was quantified in CD11b^+^ microglia using the PE channel.

### Quantitative real-time PCR (RT-qPCR)

Postnatal pups were anesthetized by intraperitoneal injection of ketamine/xylazine and decapitated. Hippocampus and cortex tissue were dissected and snap-frozen in liquid nitrogen. Total RNA was extracted using TRIzol Reagent (Invitrogen, 15596026) and used for cDNA synthesis with RNA to cDNA EcoDry Premix (Oligo dT, TaKaRa, 639541). Quantitative real-time polymerase chain reaction (RT-qPCR) was performed using AzuraView™ GreenFast qPCR Blue Mix LR (Azura Genomics, AZ-2301) on a Rotor-Gene Q thermocycler (Qiagen). Primer sequences used for qPCR analysis were as follows: *P2ry12*, forward 5’-cattgaccgctacctgaagacc-3’ and reverse 5’-gcctcctgttggtgagaatcatg-3’; *Tnfα*, forward 5’-ttgtctactcccaggttctct-3’ and reverse 5’-gaggttgactttctcctggtatg-3’; *Il6*, forward 5’-acacatgttctctgggaaatc-3’ and reverse 5’-aagtgcatcatcgttgttcatac-3’; *Il1β*, forward 5’-ggtgtgtgacgttcccatta-3’ and reverse 5’-attgaggtggagagctttcag-3’; *Cyclo* (*Cyclophilin B*), forward 5 ‘-tggagagcaccaagacagaca-3’ and reverse 5’-tgccggagtcgacaatgat-3’ (26,27). *Cyclo* was used as the housekeeping gene and relative mRNA expression levels were calculated using the 2^-ΔΔCt^ method.

### Preparation of Sindbis virus

Recombinant Sindbis viruses were generated using CT84, CT100, and helper plasmids as previously described (28–30). Viral RNA was synthesized from linearized plasmids using the mMESSAGE mMACHINE^TM^ SP6 Transcription kit (Invitrogen, AM1340) and electroporated into BHK-21 cells using the Cell Line Nucleofector kit (Lonza, VCA-1005) with a Nucleofector^TM^ I device (Amaxa). Cells were incubated for 36 h, the culture supernatant was collected, and cell debris was removed before centrifugation at 150,000 × g for 90 min at 4 °C. The virus pellet was resuspended in 200 μl BHK-21 cell culture medium consisting of MEM supplemented with 10% regular fetal bovine serum and 50 μg/ml gentamicin, aliquoted and stored at −80 °C until use. Viral stocks were diluted 1:30 for sparse neuronal infection. Neurons infected with CT84 or CT100 expressed tdTomato.

### Preparation of organotypic hippocampal slice cultures

Organotypic hippocampal slice cultures (OHSCs) were prepared from postnatal day 7 (P7) pups as described (28,31). Hippocampi were sectioned at a thickness of 250 μm using a Stoelting^TM^ tissue slicer. Slices were transferred onto Millicell^®^ six-well culture inserts (Millipore, PICM0RG50) and maintained at 37 °C in a humidified incubator with 5% CO_2_. The slice culture medium consisted of 50% MEM (Gibco, 12360038), 25% EBSS (Gibco, 24010043), 25% heat-inactivated horse serum, 2 mM GlutaMAX (Gibco, 35050061), 1 mg/l insulin, 0.00125% ascorbic acid, and 50 μg/ml gentamicin (Gibco, 15710064). The culture medium was replaced every 2 days.

Slices obtained from each mouse brain were randomly distributed between the CT84 and CT100 infection groups to minimize intra-animal variability. Slice cultures were maintained for 7 days *in vitro* before Sindbis virus infection. Virus was microinjected into the CA1 region using a FemtoJet microinjector (Eppendorf) mounted on a mechanical micromanipulator (Narishige) with pre-pulled microcapillary needles (Tritech Research, MINJ-PP without internal filament). Recombinant proteins were allowed to express for 18 h before fixation.

### Analysis of dendritic spine density

Imaging was acquired using a Nikon AXR confocal microscope with a 100× oil-immersion objective and 3.8× digital zoom. The CA1 region was identified by DAPI staining, and pyramidal neurons were identified by morphology of tdTomato^+^ cells. Dendritic segments approximately 100–200 μm from the neuron soma were imaged using Z-stack acquisition at 0.3 μm intervals with a 1024 × 1024-pixel resolution.

Spine density analysis was conducted as previously described (28,29). Briefly, dendrite length and total spine number were manually quantified using NIS-Elements Viewer (Nikon) and ImageJ (NIH) for dendritic segments with a diameter >1 μm. Values from multiple segments belonging to the same neuron were averaged to yield a single data point per neuron, expressed as the number of spines per 10 μm of dendrite.

### Reanalysis of published bulk RNA-seq data

Raw gene count data from isolated spinal microglia of adult control and ABC-imKD mice were obtained from the Gene Expression Omnibus (GEO; accession GSE154816) (17). The dataset included three biological replicates (mice) per genotype. Raw gene counts were analyzed using the DESeq2 package in R and subjected to variance-stabilizing transformation (VST). For heatmap visualization, VST-transformed expression values for selected cholesterol transport-related genes (*Abca1* and *Abcg1*) and oxidative stress-related genes (*Cybb*, *Ncf1*, *Hmox1*, and *Gclc*) were Z-score transformed across samples for each gene. Technical replicates are displayed individually in the heatmap.

### Statistical analysis

Data are presented as mean ± standard error of the mean (SEM). Statistical outliers were identified and excluded using the ROUT method (Q = 1%) in GraphPad Prism 10. For flow cytometry, histograms are normalized to the mode of each sample to allow comparison of relative frequencies regardless of differences in total event counts. Statistical significance between two groups was determined using two-tailed unpaired *t*-tests with Welch’s correction. For comparisons among four groups, data were assessed for normality and tested for homogeneity of variance using the Brown-Forsythe test. When data failed the normality test (α = 0.05), differences were analyzed using a Kruskal–Wallis test followed by Dunn’s multiple comparisons test. Normal data with homogeneous variances were analyzed using ordinary one-way ANOVA with Tukey’s multiple comparisons test. When variance homogeneity was violated, data were analyzed using Brown-Forsythe and Welch ANOVA tests with Dunnett’s T3 multiple comparisons test. Statistical details are provided in the corresponding figure legends.

## Results

### Microglial ABCA1/ABCG1 deficiency leads to cholesterol accumulation

To delete the cholesterol transporters ABCA1 and ABCG1 in microglia, we used a conditional, TAM-inducible Cre-loxP *Abca1^fl/fl^ Abcg1^fl/fl^ Cx3cr1-Cre^ERT2^* mouse model (ABC-imKD) (17). *Abca1^fl/fl^ Abcg1^fl/fl^* mice lacking *Cx3cr1-Cre^ERT2^* allele were used as controls (WT). TAM or corn oil (vehicle, Veh) was administered daily from postnatal day 1 to 3, and the efficiency of ABCA1/ABCG1 depletion was assessed at approximately postnatal day 7 (P7) using immunohistochemistry and flow cytometry (**Fig. 1A**). Immunohistochemical analysis showed that ABCA1 protein levels in IBA1^+^ microglia were reduced by an average of 34% in TAM-treated ABC-imKD mice compared with that in vehicle-treated ABC-imKD mice (**Fig. 1B**). In contrast, microglial ABCA1 expression did not differ significantly between TAM- and vehicle-treated WT mice (**Fig. S1A**). We also observed a modest trend toward reduction in ABCG1 expression in IBA1^+^ microglia from TAM-treated ABC-imKD mice compared with that in vehicle-treated ABC-imKD mice, with an average reduction of 14% (**Fig. S1B**), which may reflect the limited sensitivity or specificity of the ABCG1 antibody. Consistent with the ABCA1 results, ABCG1 expression in microglia did not differ significantly between TAM- and vehicle-treated WT mice (**Fig. S1C**). In addition to assessing ABCA1 expression *in situ* by immunohistochemistry, we used microglia-enriched brain cell suspensions and CD11b^+^ gating to further evaluate gene knockout efficiency by flow cytometry. ABCA1 protein levels were reduced by an average of 69% in microglia from TAM-treated ABC-imKD mice compared with that in vehicle-treated ABC-imKD mice (**Fig. 1C** and **S2A**).

**Figure 1.**
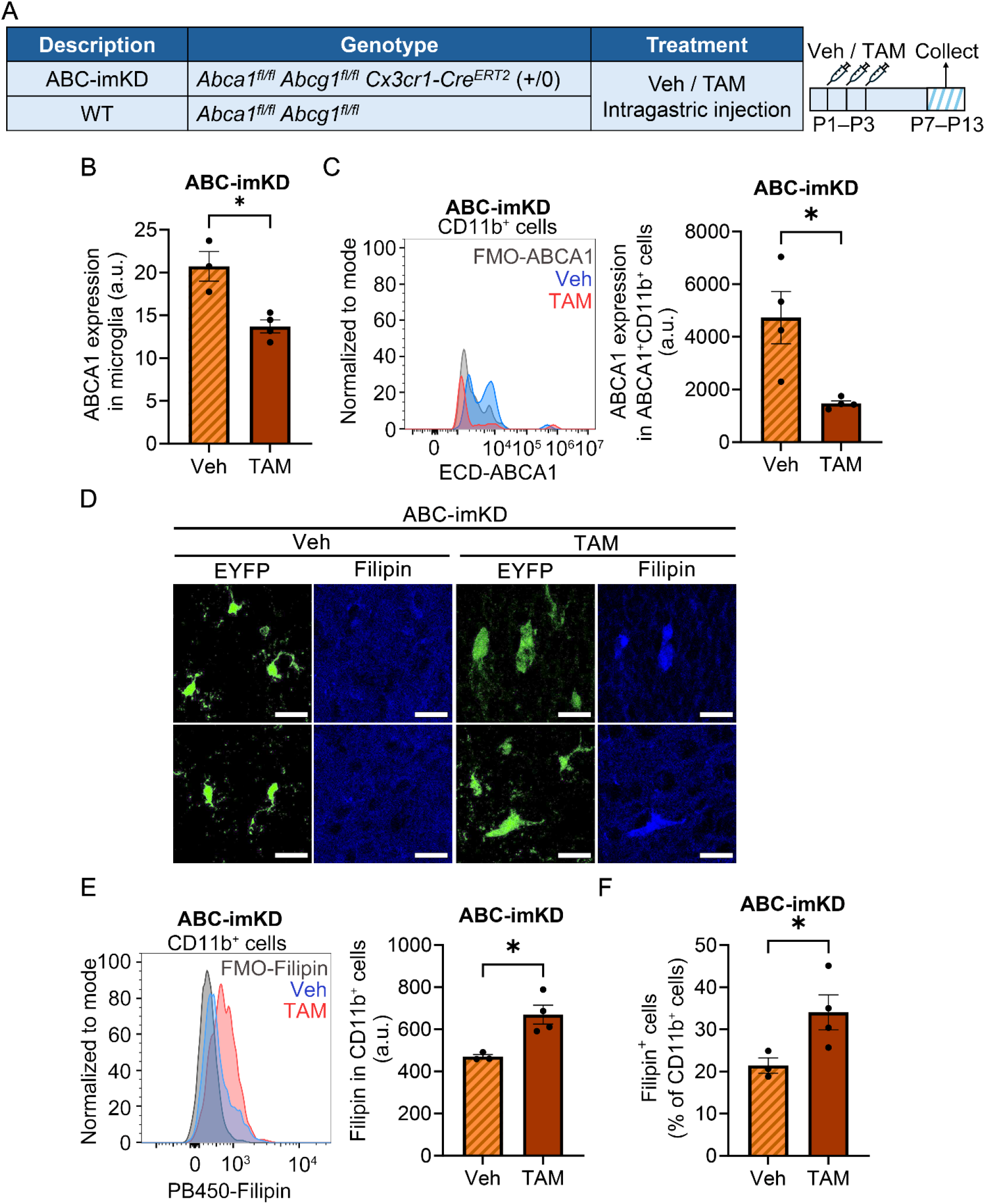
Cholesterol accumulation in microglia in mice with inducible microglial ABCA1/ABCG1 deficiency. **A,** Experimental design and genotypes of mice. ABC-imKD mice [*Abca1^fl/fl^ Abcg1^fl/fl^ Cx3cr1-Cre^ERT2^* (+/0)] and WT mice (*Abca1^fl/fl^ Abcg1^fl/fl^*) received intragastric injections of vehicle (Veh) or tamoxifen (TAM) daily from postnatal day 1 to day 3 (P1–P3). Tissues were collected at P7–P13 for immunohistochemistry or flow cytometry. **B,** Quantitative analysis of ABCA1 expression in IBA1^+^ microglia from vehicle- and tamoxifen-treated ABC-imKD mice (Veh, *n* = 3; TAM, *n* = 4). **C,** Flow cytometry analysis of ABCA1 expression in CD11b^+^ live singlets from vehicle- and tamoxifen-treated ABC-imKD mice (Veh, *n* = 4; TAM, *n* = 4). Left, representative overlay histogram of ABCA1 fluorescence in CD11b^+^ live singlets from ABC-imKD mice treated with vehicle (Veh, blue) or tamoxifen (TAM, red), with FMO-ABCA1 shown in gray, normalized to mode (% Max). FMO-ABCA1, fluorescence-minus-one control for ABCA1. ABCA1 fluorescence was detected in the ECD channel. Right, median ABCA1 fluorescence intensity in ABCA1^+^CD11b^+^ live singlets. **D,** Representative images of EYFP^+^ microglia (green) and Filipin staining (blue) in the brain of vehicle- and tamoxifen-treated ABC-imKD mice at P13. Scale bar, 20 μm. **E,** Flow cytometry analysis of Filipin fluorescence in CD11b^+^ live singlets from vehicle- and tamoxifen-treated ABC-imKD mice (Veh, *n* = 3; TAM, *n* = 4). Left, representative overlay histogram of Filipin fluorescence in CD11b^+^ live singlets from ABC-imKD mice treated with vehicle (Veh, blue) or tamoxifen (TAM, red), with FMO-Filipin shown in gray, normalized to mode (% Max). FMO-Filipin, fluorescence-minus-one control for Filipin. Filipin fluorescence was detected in the PB450 channel. Right, median Filipin fluorescence intensity in CD11b^+^ live singlets. **F,** Flow cytometry analysis of the percentage of Filipin^+^ cells among CD11b^+^ live singlets from P13 ABC-imKD mice treated with vehicle or tamoxifen (Veh, *n* = 3; TAM, *n* = 4). Statistical analyses in **B, C, E,** and **F** were performed using an unpaired two-tailed *t*-test with Welch’s correction. *, p<0.05. Data are presented as mean ± SEM. SEM, standard error of the mean. *n*, number of mice.

ATP-binding cassette transporters ABCA1/ABCG1 play important roles in regulating cellular cholesterol efflux (8). To directly assess the functional consequences of ABCA1/ABCG1 deficiency in microglia, we measured unesterified (free) cholesterol levels using Filipin staining by immunohistochemistry and flow cytometry. Because ABC-imKD mice express endogenous EYFP under the control of the *Cx3cr1* promoter, EYFP was used to identify genetically targeted microglia. Immunohistochemical analysis showed increased Filipin signal in EYFP^+^ microglia from tamoxifen-treated ABC-imKD mice, indicating cellular cholesterol accumulation (**Fig. 1D**). In contrast, Filipin signal was not different between EYFP^+^ and EYFP^-^ cells in vehicle-treated ABC-imKD mice (**Fig. 1D**). Consistent with these findings, flow cytometry analysis showed that the Filipin fluorescence intensity of CD11b^+^ microglia was increased by an average of 1.4-fold in TAM-treated ABC-imKD mice compared with that in vehicle-treated mice (**Fig. 1E** and **S2B**). In addition, flow cytometry analysis revealed a marked increase in the percentage of Filipin^+^ cells among CD11b^+^ microglia in tamoxifen-treated ABC-imKD mice compared with that in vehicle-treated mice (**Fig. 1F**). The percentage of Filipin^+^ microglia increased from an average of 21% in vehicle-treated mice to 34% in TAM-treated mice, corresponding to a 1.6-fold increase. Taken together, these results show that TAM-induced recombination reduces microglial ABCA1/ABCG1 expression and increases cellular cholesterol accumulation.

### Microglial ABCA1/ABCG1 deficiency results in a reactive microglial phenotype and increases oxidative stress

In Figure 1D, we noticed a morphological difference in EYFP^+^ microglia. EYFP^+^ microglia in the vehicle-treated group appeared ramified, with long and thin processes, whereas those in the TAM-treated group appeared more amoeboid, with enlarged and rounded cell bodies. We hypothesized that cholesterol transporter-deficient microglia underwent a phenotypic transition from the homeostatic to a reactive state. To test this hypothesis, we analyzed the morphology of IBA1^+^ microglia in the hippocampal region of TAM- and vehicle-treated ABC-imKD mice (**Fig. 2A**). TAM treatment resulted in a 1.35-fold increase in microglial cell area compared with vehicle-treated ABC-imKD mice, based on the median of IBA1^+^ cell area calculated for each mouse (**Fig. 2B**).

**Figure 2.**
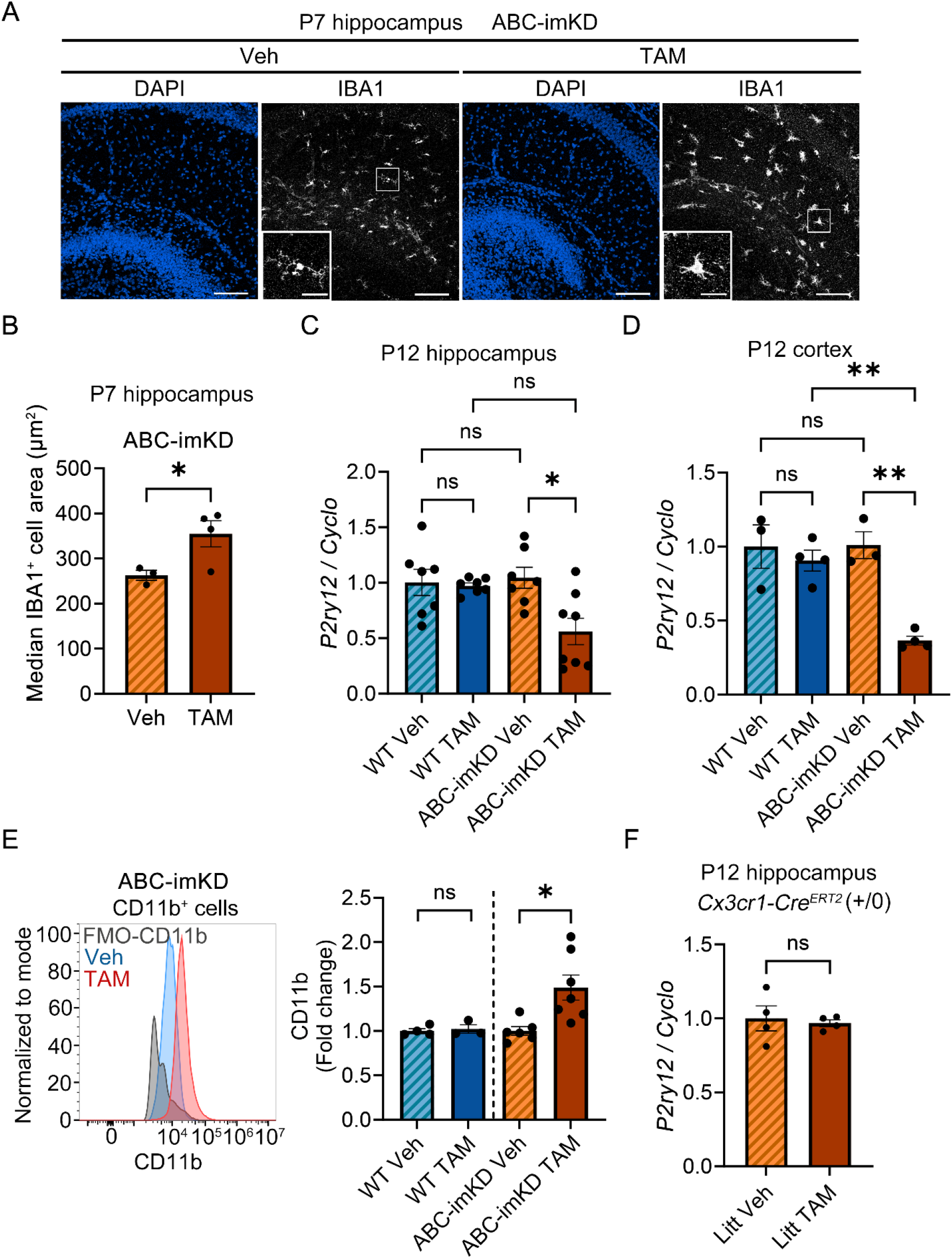
Microglial cholesterol transporter deficiency promotes a reactive microglial phenotype. **A,** Representative immunohistochemical images of the postnatal day 7 (P7) hippocampus showing DAPI (nuclear marker, blue) and IBA1 (microglial marker, white) in vehicle (Veh)- and tamoxifen (TAM)-treated ABC-imKD mice. Scale bar, 100 μm. Insets show higher-magnification views of microglia within the boxed regions. Scale bar, 25 μm. **B,** Quantification of IBA1^+^ microglial cell area in the P7 hippocampus. For each mouse, the median cell area was calculated from 199–530 individual IBA1^+^ microglia and used as a single biological replicate (Veh, *n* = 3; TAM, *n* = 4). **C,** Relative *P2ry12* mRNA expression (normalized to *Cyclo*) in the P12 hippocampus of vehicle- and tamoxifen-treated WT mice (Veh, *n* = 7; TAM, *n* = 7) and ABC-imKD mice (Veh, *n* = 7; TAM, *n* = 8). **D,** Relative *P2ry12* mRNA expression (normalized to *Cyclo*) in the P12 cortex of vehicle- and tamoxifen-treated WT mice (Veh, *n* = 3; TAM, *n* = 4) and ABC-imKD mice (Veh, *n* = 3; TAM, *n* = 4). **E,** Flow cytometry analysis of CD11b fluorescence intensity in WT mice (Veh, *n* = 4; TAM, *n* = 3) and ABC-imKD mice (Veh, *n* = 6; TAM, *n* = 7) at P11–P13. Left, representative overlay histogram of CD11b fluorescence in CD11b^+^ live singlets from vehicle- and tamoxifen-treated ABC-imKD mice (Veh, blue; TAM, red), with FMO-CD11b shown in gray, normalized to mode (% Max). FMO-CD11b, fluorescence-minus-one control for CD11b. Right, geometric mean fluorescence intensity of CD11b in CD11b^+^ live singlets. Values were normalized to the mean of the corresponding vehicle-treated group. **F,** Relative *P2ry12* mRNA expression (normalized to *Cyclo*) in the P12 hippocampus of vehicle- and tamoxifen-treated Litt mice [*Cx3cr1-Cre^ERT2^* (+/0)] (Veh, *n* = 4; TAM, *n* = 4). For **B**–**F**, ns, not significant; *, p<0.05; **, p<0.01. Data are presented as mean ± SEM. SEM, standard error of the mean. Statistical analyses were performed using an unpaired two-tailed *t*-test with Welch’s correction (**B**, **E**, and **F**), Brown–Forsythe and Welch ANOVA followed by Dunnett’s T3 multiple comparisons test (**C**), or ordinary one-way ANOVA followed by Tukey’s multiple comparisons test (**D**). *n*, number of mice.

We next isolated total RNA from the hippocampus and performed quantitative real-time polymerase chain reaction (RT-qPCR). We found that relative *P2ry12* mRNA expression was significantly decreased by an average of 46% in the P12 hippocampus of TAM-treated ABC-imKD mice compared with vehicle-treated ABC-imKD mice (**Fig. 2C**). A similar decrease in *P2ry12* expression was observed in the cortex, with an average reduction of 64% (**Fig. 2D**). These data indicate a downregulation of the microglial homeostatic marker *P2ry12* in brain tissue following microglial ABCA1/ABCG1 deficiency.

In addition to the homeostatic marker, we assessed the microglial activation-associated marker CD11b (32) by flow cytometry. CD11b fluorescence intensity in microglia was significantly increased by 1.5-fold in TAM-treated ABC-imKD mice compared with that in vehicle-treated ABC-imKD mice, whereas CD11b fluorescence intensity was not different between TAM- and vehicle-treated WT mice (**Fig. 2E**). Together, these data indicate that ABCA1/ABCG1 deficiency in microglia promotes a shift toward a reactive microglial phenotype.

Because a previous study reported that TAM-induced Cre activity in the *Cx3cr1-Cre^ERT2^* line used in our study could lead to microglial activation, with this adverse effect being reported primarily during the neonatal stage (24), we sought to exclude this potential confounding effect in our experimental system. We therefore measured *P2ry12* expression in the hippocampus of *Cx3cr1-Cre^ERT2^* mice at P12 and found no differences in *P2ry12* mRNA expression between TAM- and vehicle-treated *Cx3cr1-Cre^ERT2^* (+/0) mice (**Fig. 2F**). These results indicate that the reactive microglial phenotype observed in ABC-imKD mice is associated with ABCA1/ABCG1 deficiency rather than with TAM-induced Cre activity itself.

Primary microglia isolated from the brains of ABCA1 whole-body-deficient mice exhibited reduced lipid efflux, resulting in the accumulation of lipid droplets in *Abca1^-/-^* microglia (33). In a study of aging mouse and human brains, lipid-droplet-accumulating microglia were shown to produce high levels of reactive oxygen species (ROS) (34). We therefore asked whether reactive oxygen species were increased in cholesterol transporter-deficient microglia. Superoxide is considered an early and primary reactive oxygen species generated during oxidative stress. We used the superoxide probe dihydroethidium (DHE) to detect superoxide-associated fluorescence in our experimental system. Flow cytometry analysis showed that DHE fluorescence intensity was increased by an average of 1.5-fold in CD11b^+^ microglia in TAM-treated ABC-imKD mice compared with that in vehicle-treated ABC-imKD mice (**Fig. 3A**). We then classified CD11b^+^ microglia into DHE^+^ and DHE^-^ populations using the DHE fluorescence-minus-one (FMO) control as the gating reference. The DHE signal index was calculated as the product of the percentage of DHE^+^CD11b^+^ live singlets and the geometric mean DHE fluorescence intensity of DHE^+^CD11b^+^ live singlets (17). We found a significant 2-fold increase in the DHE signal index in TAM-treated ABC-imKD mice compared with vehicle-treated ABC-imKD mice (**Fig. 3B, 3C,** and **3D**). In contrast, the DHE signal index was not different between TAM- and vehicle-treated WT mice (**Fig. 3E**). Therefore, microglial ABCA1/ABCG1 deficiency is associated with increased superoxide-associated DHE fluorescence, consistent with enhanced oxidative stress.

**Figure 3.**
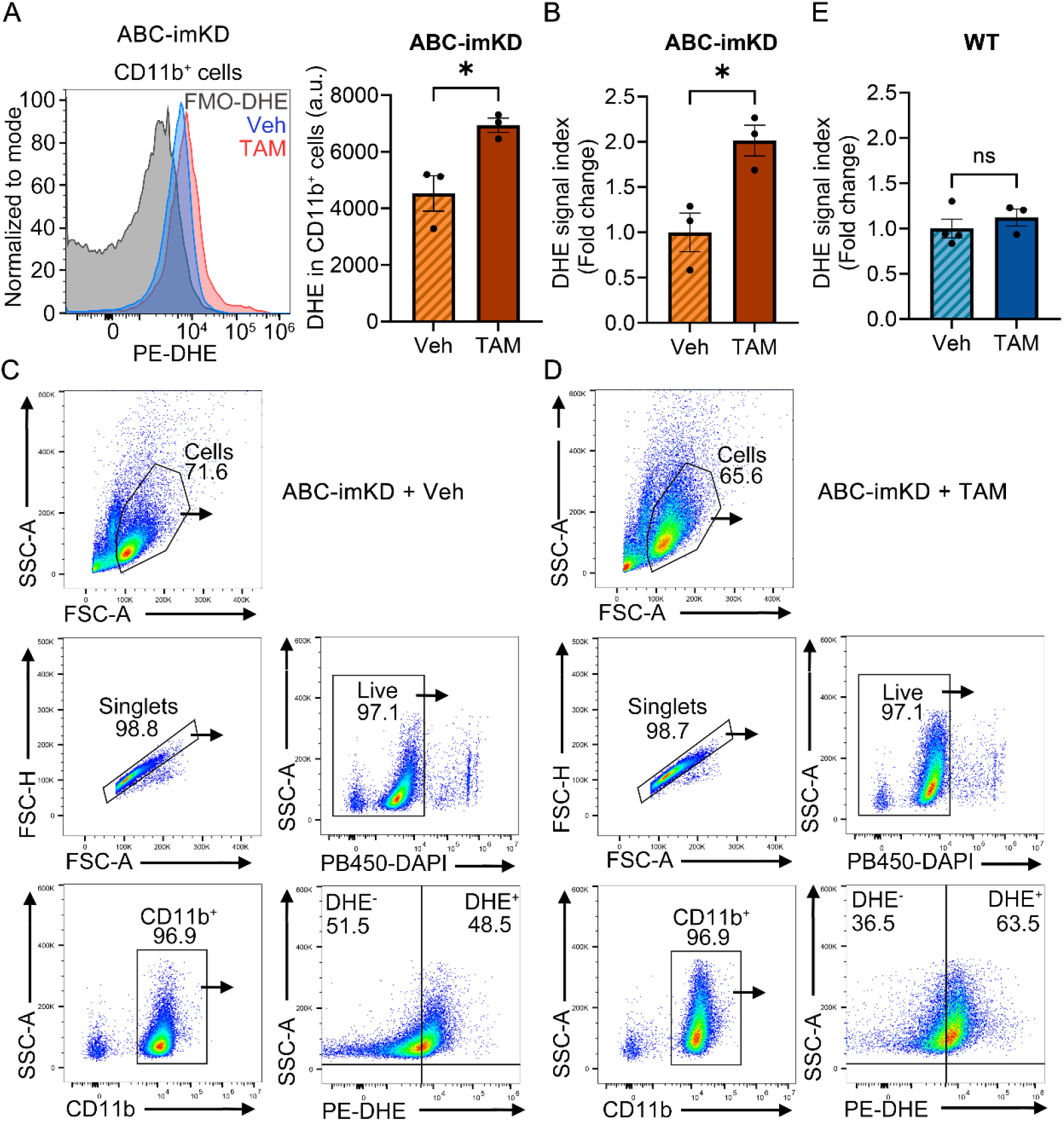
Microglial cholesterol transporter deficiency increases oxidative stress in microglia. **A,** Flow cytometry analysis of the ROS probe dihydroethidium (DHE) fluorescence in CD11b^+^ live singlets from vehicle (Veh)- and tamoxifen (TAM)-treated ABC-imKD mice (Veh, *n* = 3; TAM, *n* = 3). Left, representative overlay histogram of DHE fluorescence in CD11b^+^ live singlets from vehicle- and tamoxifen-treated ABC-imKD mice (Veh, blue; TAM, red), with FMO-DHE shown in gray, normalized to mode (% Max). FMO-DHE, fluorescence-minus-one control for DHE. Right, geometric mean DHE fluorescence intensity in CD11b^+^ live singlets. **B** and **E,** Quantification of the DHE signal index in CD11b^+^ cells from vehicle- and tamoxifen-treated ABC-imKD (**B**) and WT mice (**E**). The DHE signal index was calculated as the product of the percentage of DHE^+^CD11b^+^ live singlets and the geometric mean DHE fluorescence intensity of DHE^+^CD11b^+^ live singlets. Values were normalized to the mean of the corresponding vehicle-treated group and expressed as fold change. For ABC-imKD mice, Veh, *n* = 3; TAM, *n* = 3. For WT mice, Veh, *n* = 4; TAM, *n* = 3. **C**–**D,** Representative gating strategy for sequential identification of cells, singlets, live cells, CD11b^+^ cells, and DHE^+^CD11b^+^ cells from vehicle-treated (**C**) and tamoxifen-treated (**D**) ABC-imKD mice. For **A**, **B** and **E**, *n*, number of mice. ns, not significant; *, p<0.05. Data are presented as mean ± SEM. Statistical analyses were performed using an unpaired two-tailed *t*-test with Welch’s correction.

In addition, we reanalyzed our bulk RNA-seq data from adult ABC-imKD spinal cord microglia (17) and found that oxidative stress-related genes, including *Cybb* (NOX2), *Ncf1*, *Hmox1*, and *Gclc*, showed upward trends in ABCA1/ABCG1-deficient microglia compared with control microglia (**Fig. S3**). These results suggest that ABCA1/ABCG1 deficiency confers a transcriptional profile consistent with increased oxidative stress and possible involvement of NOX2-associated pathways.

### Microglial ABCA1/ABCG1 deficiency is associated with increased susceptibility to Aβ-induced dendritic spine loss

Having established that ABCA1/ABCG1 deficiency disrupts microglial cholesterol homeostasis and promotes a reactive phenotype associated with increased oxidative stress, we next asked whether these alterations affect neuronal synaptic integrity. To address this question, we assessed dendritic spine density in organotypic hippocampal slice cultures (OHSCs) from vehicle- and TAM-treated mice under basal and Aβ-stressed conditions (**Fig. 4A** and **4B**). Aβ-associated stress was induced using a Sindbis viral system expressing CT100, an APP C-terminal fragment containing the Aβ sequence that induces neuronal Aβ expression, whereas CT84, the corresponding fragment lacking the Aβ sequence, served as the control (28). Neurons were co-labelled with tdTomato to identify infected neurons and visualize dendritic spines in CA1 pyramidal neurons. Previous studies using this system showed that CT100 expression induced time-dependent dendritic spine loss, with a significant reduction observed at 18–24 h, but not at 12 h, after infection (29).

**Figure 4.**
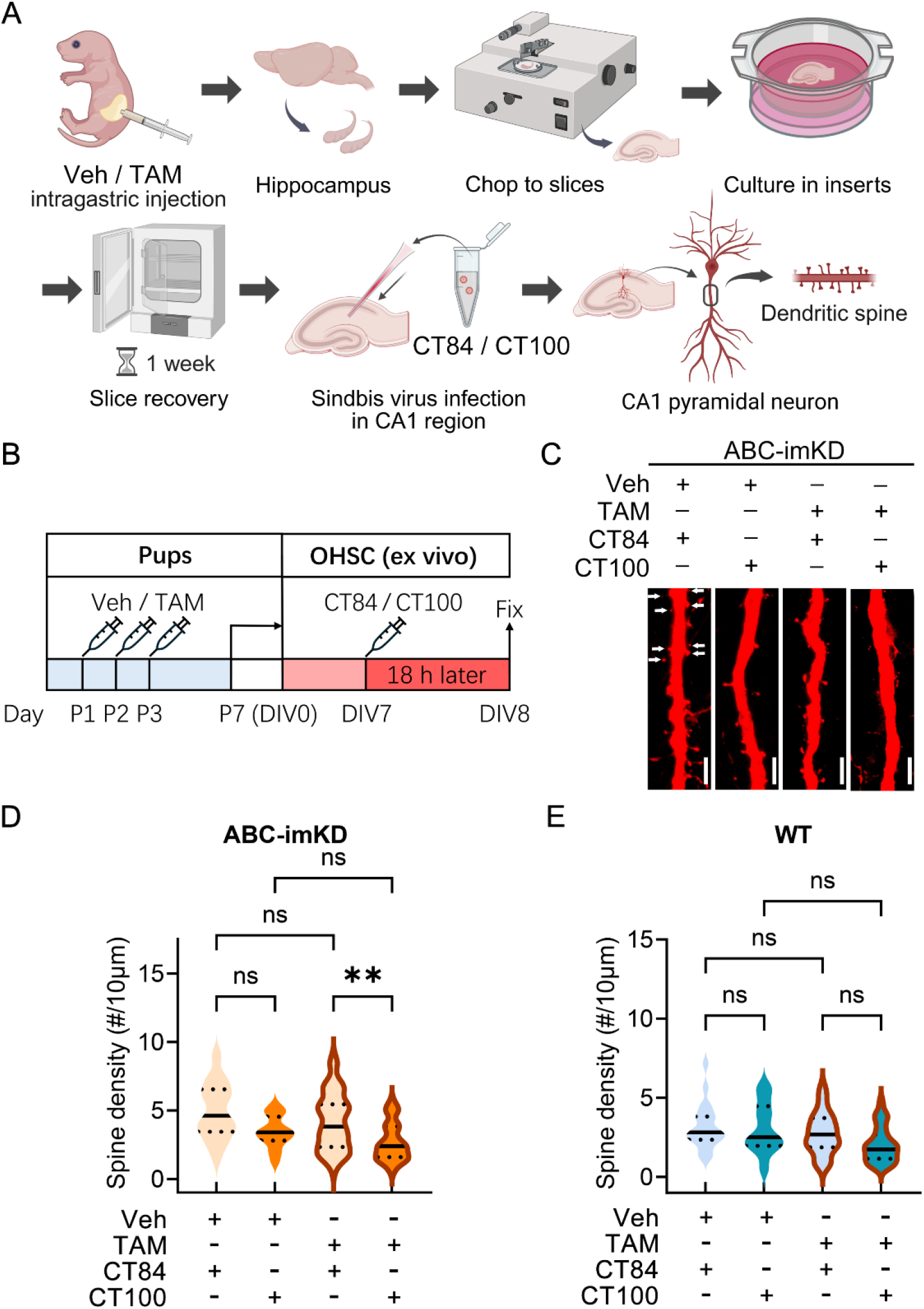
Microglial ABCA1/ABCG1 deficiency increases susceptibility to Aβ-induced dendritic spine loss. **A,** Schematic illustration of the experimental workflow for organotypic hippocampal slice culture (OHSC) and dendritic spine analysis. Hippocampi were dissected from vehicle (Veh)- or tamoxifen (TAM)-treated postnatal mice and prepared as OHSCs. Following a 1-week recovery period, slices were infected with Sindbis virus expressing CT84 or CT100 in the CA1 region, followed by dendritic spine analysis of tdTomato^+^ CA1 pyramidal neurons. Regions of interest for dendritic spine analysis were located approximately 100–200 μm from the soma. **B,** Experimental timeline. Neonatal pups received intragastric injections of tamoxifen (50 μg) or an equivalent volume of vehicle daily from postnatal day 1 to day 3 (P1–P3), and OHSCs were prepared at P7 [day *in vitro* 0 (DIV0)]. At DIV7, slice cultures were infected with Sindbis virus expressing either the control construct (CT84) or the amyloid-beta (Aβ)-generating construct (CT100) together with tdTomato. Slice cultures were fixed with 4% paraformaldehyde 18 h after infection and analyzed at DIV8. **C,** Representative images of dendritic spines from neurons expressing CT84 or CT100 in OHSCs prepared from vehicle- and tamoxifen-treated ABC-imKD mice. White arrows indicate dendritic spines. Scale bar, 5 μm. **D,** Quantification of dendritic spine density in OHSCs from vehicle- and tamoxifen-treated ABC-imKD mice infected with Sindbis virus expressing CT84 or CT100. *n* = 29 neurons (Veh, CT84), *n* = 24 neurons (Veh, CT100), *n* = 37 neurons (TAM, CT84), and *n* = 47 neurons (TAM, CT100), obtained from 4–5 ABC-imKD mice per group. **E,** Quantification of dendritic spine density in OHSCs from vehicle- and tamoxifen-treated WT mice infected with Sindbis virus expressing CT84 or CT100. *n* = 19 neurons (Veh, CT84), *n* = 20 neurons (Veh, CT100), *n* = 21 neurons (TAM, CT84), and *n* = 33 neurons (TAM, CT100), obtained from 5–6 WT mice per group. For the violin plots in **D** and **E**, solid lines indicate the median, and dotted lines indicate the first and third quartiles. Light colors (light orange in **D** and light cyan in **E**) represent CT84 infection, whereas dark colors (dark orange in **D** and dark cyan in **E**) represent CT100 infection. Violin plots without brown outlines represent vehicle-treated mice, whereas violin plots with brown outlines represent tamoxifen-treated mice. Statistical analyses were performed using the Kruskal–Wallis test followed by Dunn’s multiple comparisons test. ns, not significant; **, p<0.01. *n*, number of neurons.

In ABC-imKD mice (**Fig. 4C** and **4D**), the spine density of CT84-infected CA1 pyramidal neurons 18 h post-infection was not significantly different between TAM- and vehicle-treated mice, indicating that microglial ABCA1/ABCG1 deficiency did not significantly affect basal spine density. In TAM-treated ABC-imKD mice, CT100 infection significantly reduced spine density relative to CT84 infection, with a median decrease of 35%. In contrast, CT100 infection produced a smaller median decrease of 11% in vehicle-treated ABC-imKD mice, which did not reach statistical significance. Thus, CT100-induced dendritic spine loss was more pronounced in the setting of microglial ABCA1/ABCG1 deficiency, indicating increased susceptibility to Aβ-associated synaptic stress.

We also performed the same experiments in WT mice. No significant differences in spine density were observed between CT100- and CT84-infected neurons at 18 h in either TAM- or vehicle-treated WT mice, nor was there a significant difference between TAM- and vehicle-treated mice among CT84-infected neurons (**Fig. 4E**).

Although modest reductions in dendritic spine density were observed in the control conditions, these changes did not reach statistical significance, indicating that neither TAM exposure alone nor *Cx3cr1-Cre^ERT2^* expression alone was sufficient to produce a significant Aβ-induced spine loss under these experimental conditions. Taken together, these findings demonstrate that microglial ABCA1/ABCG1 deficiency increases neuronal susceptibility to Aβ-induced dendritic spine loss.

## Discussion

Dysregulated lipid metabolism is an important feature of AD, with altered cholesterol homeostasis and the strong association of APOE with AD highlighting the importance of lipid transport in disease pathogenesis (35,36). However, how cholesterol homeostasis specifically in microglia influences neuronal integrity and the response to Aβ-associated stress remains poorly understood. Here, we found that microglial ABCA1/ABCG1 deficiency caused excessive cholesterol accumulation, a reactive microglial phenotype, and increased oxidative stress. Although these changes did not significantly affect basal dendritic spine density, they were associated with markedly greater spine loss following Aβ-associated stress (**Fig. 5**). Thus, our findings link impaired microglial cholesterol homeostasis to increased synaptic vulnerability to Aβ and identify microglial lipid metabolism as an important component of neuronal resilience under AD-related stress.

**Figure 5.**
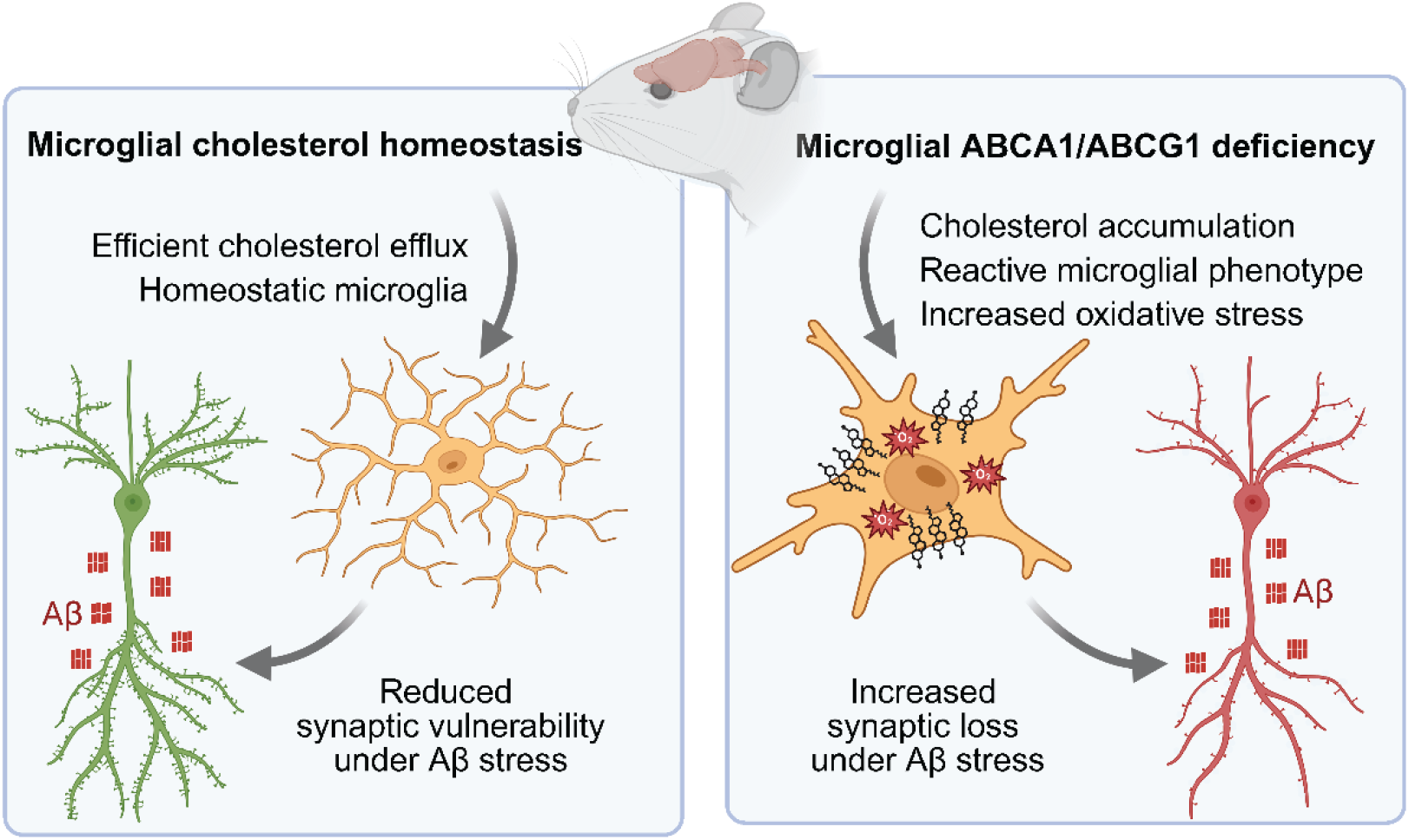
Working model of cholesterol-dependent reactive microglia phenotype and synaptic vulnerability to Aβ. Microglial ABCA1/ABCG1 maintain cholesterol homeostasis by promoting cholesterol efflux, thereby limiting microglial reactivity and oxidative stress. Under amyloid-beta (Aβ) stress, this homeostatic state helps limit synaptic vulnerability. In contrast, microglial ABCA1/ABCG1 deficiency disrupts cholesterol homeostasis, leading to cholesterol accumulation, a reactive microglial phenotype, and increased oxidative stress. These changes are proposed to enhance synaptic vulnerability to Aβ and promote dendritic spine loss.

Loss of ABCA1/ABCG1 resulted in marked cholesterol accumulation accompanied by morphological changes, increased CD11b expression, and reduced *P2ry12*, indicating that impaired cholesterol efflux alters the microglial state. P2RY12 is associated with homeostatic microglial function and is downregulated following inflammatory activation (37); thus, reduced *P2ry12* in ABCA1/ABCG1-deficient microglia is consistent with a shift away from a homeostatic phenotype. However, this altered microglial phenotype was not accompanied by significant increases in *Tnfα*, *Il6*, or *Il1β* expression in the hippocampus or cortex (**Fig. S4A**–**F**), suggesting that microglial ABCA1/ABCG1 deficiency alone does not induce a robust tissue-level pro-inflammatory cytokine response under basal conditions.

Previous evidence suggests that the inflammatory consequences of ABCA1 deficiency are context dependent and may vary according to the affected CNS cell population and the presence of an inflammatory challenge (14). Karasinska et al. reported increased mRNA expression of several inflammation-associated markers, including TNFα, iNOS, NFκB, and TGFβ, together with cortical astrogliosis in young mice with broad brain ABCA1 deficiency, whereas neuron-specific ABCA1 deletion induced astrogliosis without increased inflammatory gene expression and astrocyte-specific deletion produced neither phenotype. Importantly, primary microglia isolated from mice with broad, nestin-driven ABCA1 deficiency exhibited an exaggerated pro-inflammatory response to lipopolysaccharide (LPS), including increased TNFα production, despite only a partial (35%) reduction in microglial ABCA1 protein. Because ABCA1 was broadly reduced throughout the CNS in this model, these experiments cannot distinguish microglia-intrinsic effects of ABCA1 deficiency from prior influences of the ABCA1-deficient brain environment. Nevertheless, the enhanced response of isolated microglia to LPS is consistent with the possibility that impaired ABCA1-dependent cholesterol homeostasis places microglia in a sensitized or primed state, in which inflammatory responses become more apparent following a subsequent challenge rather than being constitutively elevated under basal conditions. This concept is consistent with our observation that ABCA1/ABCG1-deficient microglia exhibit an altered phenotype and increased oxidative stress at baseline, whereas neuronal consequences become particularly evident in the presence of Aβ-associated stress.

Despite the lack of a robust basal tissue-level cytokine response, ABCA1/ABCG1-deficient microglia exhibited increased superoxide levels together with increased CD11b expression. Notably, CD11b has been linked to microglial ROS production and neuronal injury in multiple contexts. In the developing hippocampus, CD11b, together with DAP12, contributes to microglial superoxide production and microglia-mediated neuronal apoptosis (32), while in rotenone-treated microglia, CD11b promotes NOX2 activation and ROS production, and disruption of the CD11b– NOX2 axis attenuates microglial activation and neurotoxicity (38). These observations raise the possibility that increased CD11b expression in ABCA1/ABCG1-deficient microglia may be associated with enhanced oxidative stress and neuronal vulnerability, potentially involving NOX2-related mechanisms. Consistent with this possibility, in our reanalysis of RNA-seq data from adult spinal microglia, the NOX2-associated genes *Cybb* and *Ncf1*, together with the oxidative stress-responsive genes *Hmox1* and *Gclc*, showed upward trends in ABCA1/ABCG1-deficient microglia, although these changes did not reach statistical significance (**Fig. S3**). Together, these findings are consistent with increased oxidative stress and possible involvement of NOX2-associated pathways. The selective exacerbation of dendritic spine loss under Aβ stress further suggests that impaired microglial cholesterol homeostasis increases synaptic vulnerability rather than causing substantial basal synaptic injury. Because oxidative stress has been linked to synaptic dysfunction in AD (39), increased microglial ROS may represent one potential link between microglial cholesterol accumulation and enhanced Aβ-induced spine loss. However, whether ROS directly contributes to this synaptic phenotype remains to be determined.

Previous studies have shown that CT100 expression in organotypic hippocampal slices induces synaptic dysfunction and time-dependent dendritic spine loss (28,29,40,41). In our study, however, CT100 did not significantly reduce spine density in WT slices at 18 h post-infection (**Fig. 4E**). This discrepancy may reflect differences in experimental timing, as slices in our studies were infected earlier, at 7 days in vitro, whereas previous studies performed infection at 13–15 days in vitro and generally allowed longer CT100 expression for 20–30 h (28–30,40). Thus, the earlier infection time point and shorter CT100 expression period may have limited the extent of CT100-induced spine loss in WT neurons.Several limitations should be considered when interpreting these findings. Although increased superoxide levels and changes in NOX2-associated oxidative pathways suggest a potential role for oxidative stress, its causal contribution to Aβ-induced spine loss remains to be established through ROS inhibition or rescue experiments. In addition, inflammatory cytokines were measured in whole hippocampal and cortical tissues under basal conditions, which may not fully capture microglia-specific inflammatory changes. Given previous evidence that primary microglia from mice with broad CNS ABCA1 deficiency can exhibit enhanced inflammatory responses to LPS, whether ABCA1/ABCG1-deficient microglia exhibit exaggerated inflammatory responses to Aβ or LPS in our model remains to be determined. Finally, CX3CR1-lineage targeting does not completely exclude contributions from infiltrating macrophages, and additional, more microglia-restricted genetic models will therefore be important to independently confirm the contribution of microglial ABCA1/ABCG1 deficiency to the observed phenotype. Despite these limitations, our findings demonstrate that disruption of ABCA1/ABCG1-dependent cholesterol homeostasis in microglia alters their phenotype and increases neuronal susceptibility to Aβ-induced dendritic spine loss, supporting further investigation of microglial cholesterol efflux as a potential target for preserving synaptic resilience to Aβ-associated stress.

## Acknowledgments

We thank Yixing Du and Amber Lawrence in Dr. Kim Dore’s lab for technical guidance and expertise. We thank Dr. Nicholas Webster (UC San Diego) for generously providing access to a flow cytometer in his laboratory. Microscopy was performed at the Nikon Imaging Center and Microscopy Core at UC San Diego. We thank both facilities for their support with image acquisition and analysis.

## Author Contributions

Y.I.M. and S.D. conceived the project, Y.I.M., S.D., and S.-H.C. designed the experiments, S.D. and N.N. performed experiments, S.-H.C. contributed to study discussions, J.K. provided the mouse breeders, K.D. provided technical guidance and expertise for the OHSC experiments, S.D. and Y.I.M. wrote the manuscript.

---

**Figure S1.**
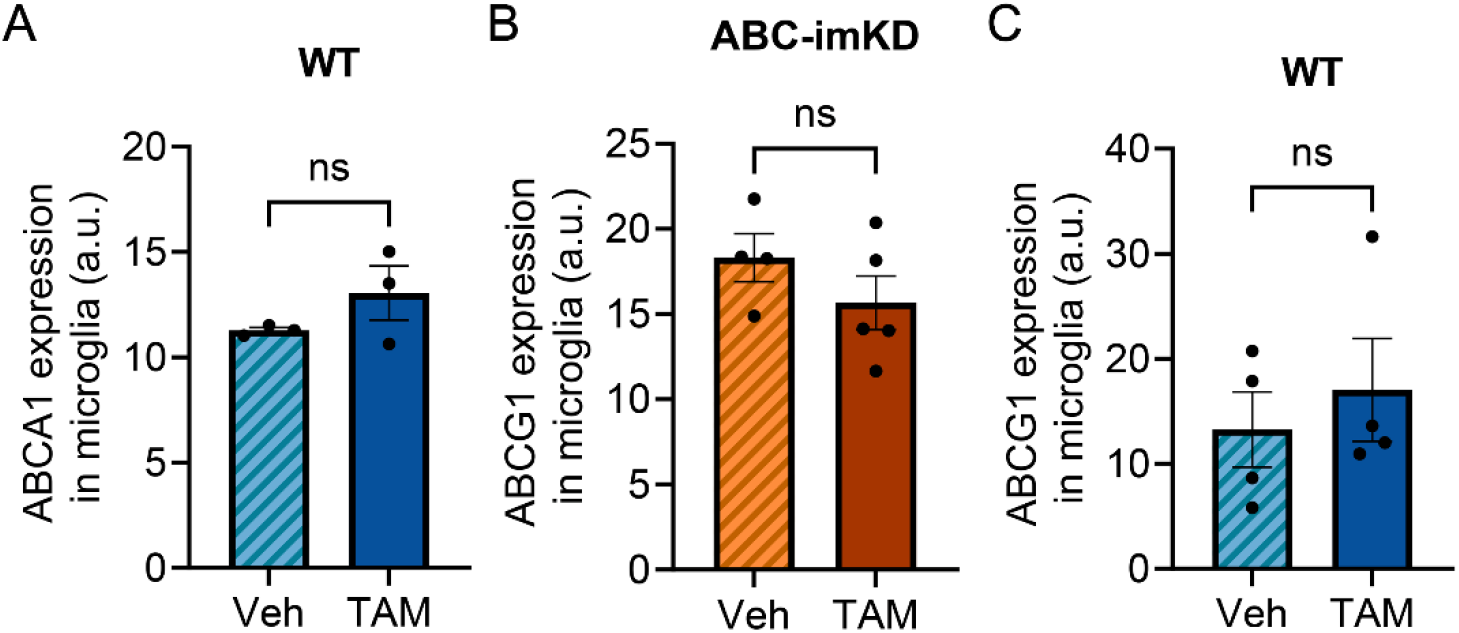
Microglial ABCA1 and ABCG1 expression in P7 WT and ABC-imKD mice. **A,** Quantitative analysis of postnatal day 7 (P7) hippocampal microglial ABCA1 expression in vehicle-treated (Veh, *n* = 3) and tamoxifen-treated (TAM, *n* = 3) WT mice. **B,** Quantitative analysis of hippocampal microglial ABCG1 expression in vehicle- and tamoxifen-treated ABC-imKD mice (Veh, *n* = 4; TAM, *n* = 5). **C,** Quantitative analysis of hippocampal microglial ABCG1 expression in vehicle- and tamoxifen-treated WT mice (Veh, *n* = 4; TAM, *n* = 4). For **A**–**C**, ns, not significant. Data are presented as mean ± SEM. SEM, standard error of the mean. *n*, number of mice. Statistical analyses were performed using an unpaired two-tailed *t*-test with Welch’s correction.

**Figure S2.**
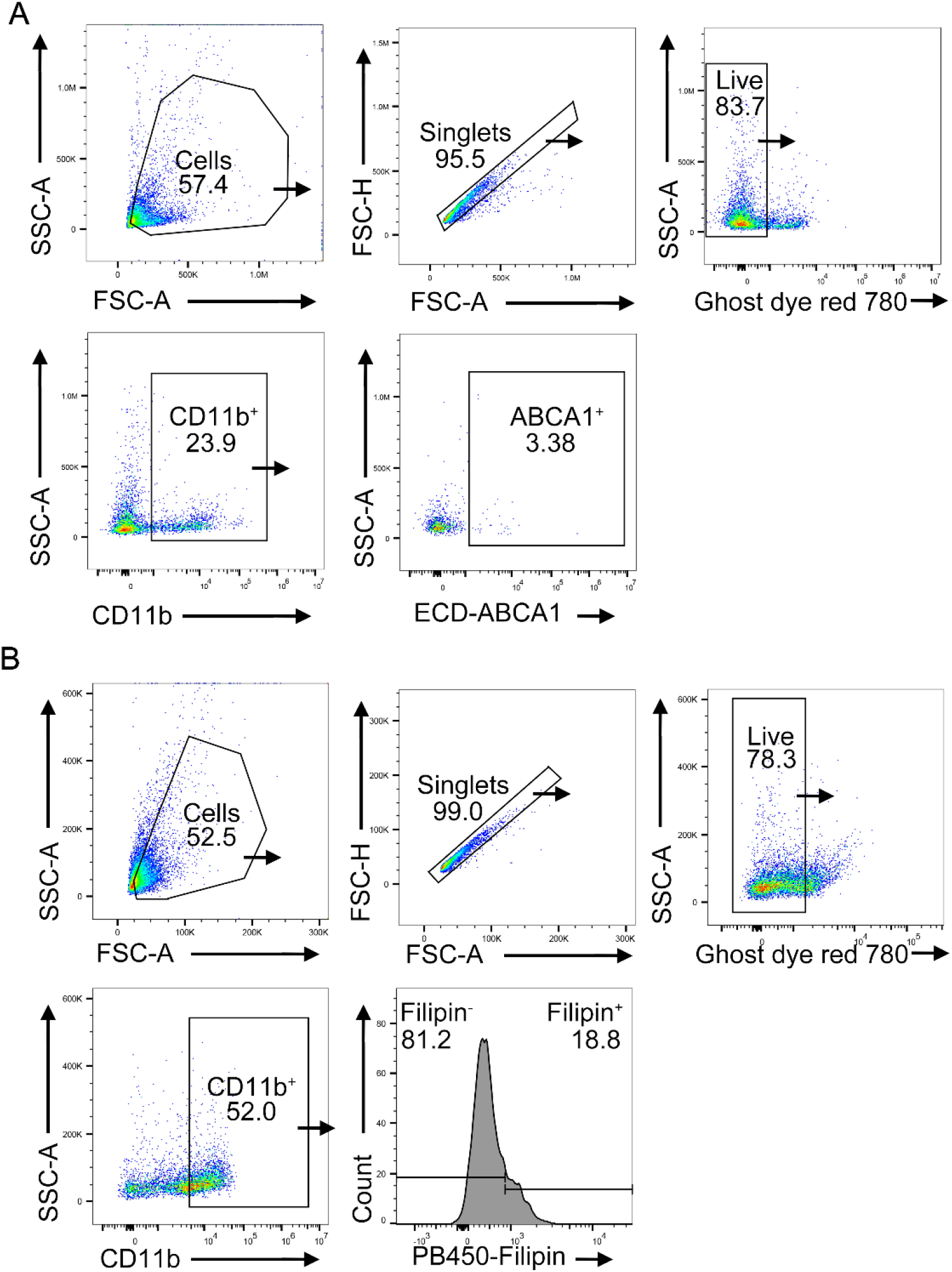
Flow cytometry gating strategy for analysis of microglial ABCA1 expression and Filipin staining. **A,** Flow cytometry gating strategy and representative plots of ABCA1 expression in CD11b^+^ live singlets. ABCA1 expression was detected in the ECD channel. **B,** Flow cytometry gating strategy and representative plots of Filipin staining in CD11b^+^ live singlets. Filipin staining was detected in the PB450 channel.

**Figure S3.**
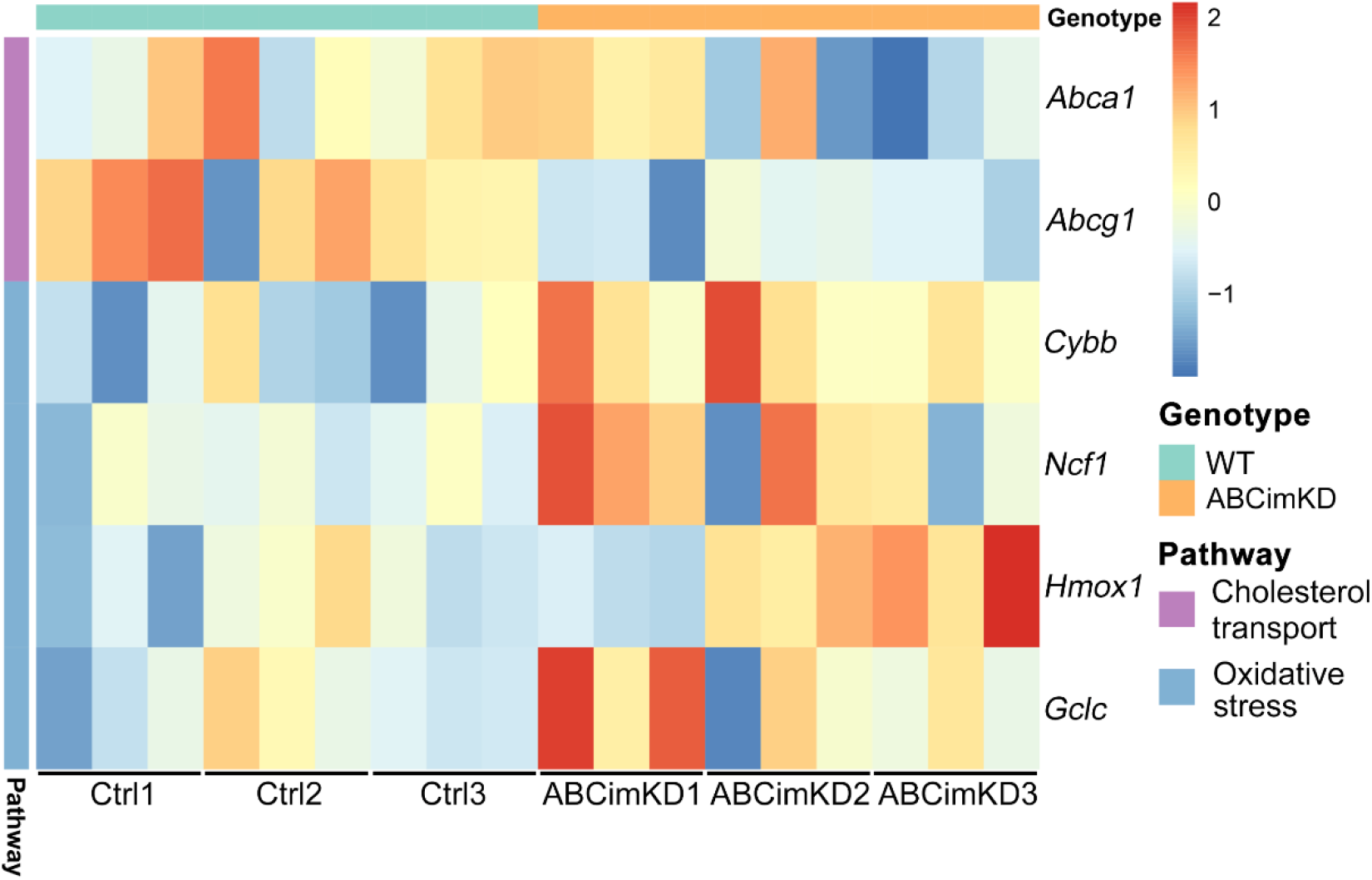
Expression patterns of selected cholesterol transport- and oxidative stress-related genes in adult control and ABC-imKD spinal microglia. Reanalysis of our published bulk RNA-seq data from isolated spinal microglia of adult ABC-imKD (ABCimKD) and control (Ctrl; C57BL/6J background wild-type) mice showing expression patterns of the cholesterol transporters *Abca1* and *Abcg1* and oxidative stress-related genes *Cybb*, *Ncf1*, *Hmox1*, and *Gclc*. Each genotype included three biological replicates (mice), with three technical replicates per mouse. Heatmap colors represent Z-scores calculated across samples for each gene after variance-stabilizing transformation. Column and row annotations indicate genotype and functional gene category, respectively.

**Figure S4.**
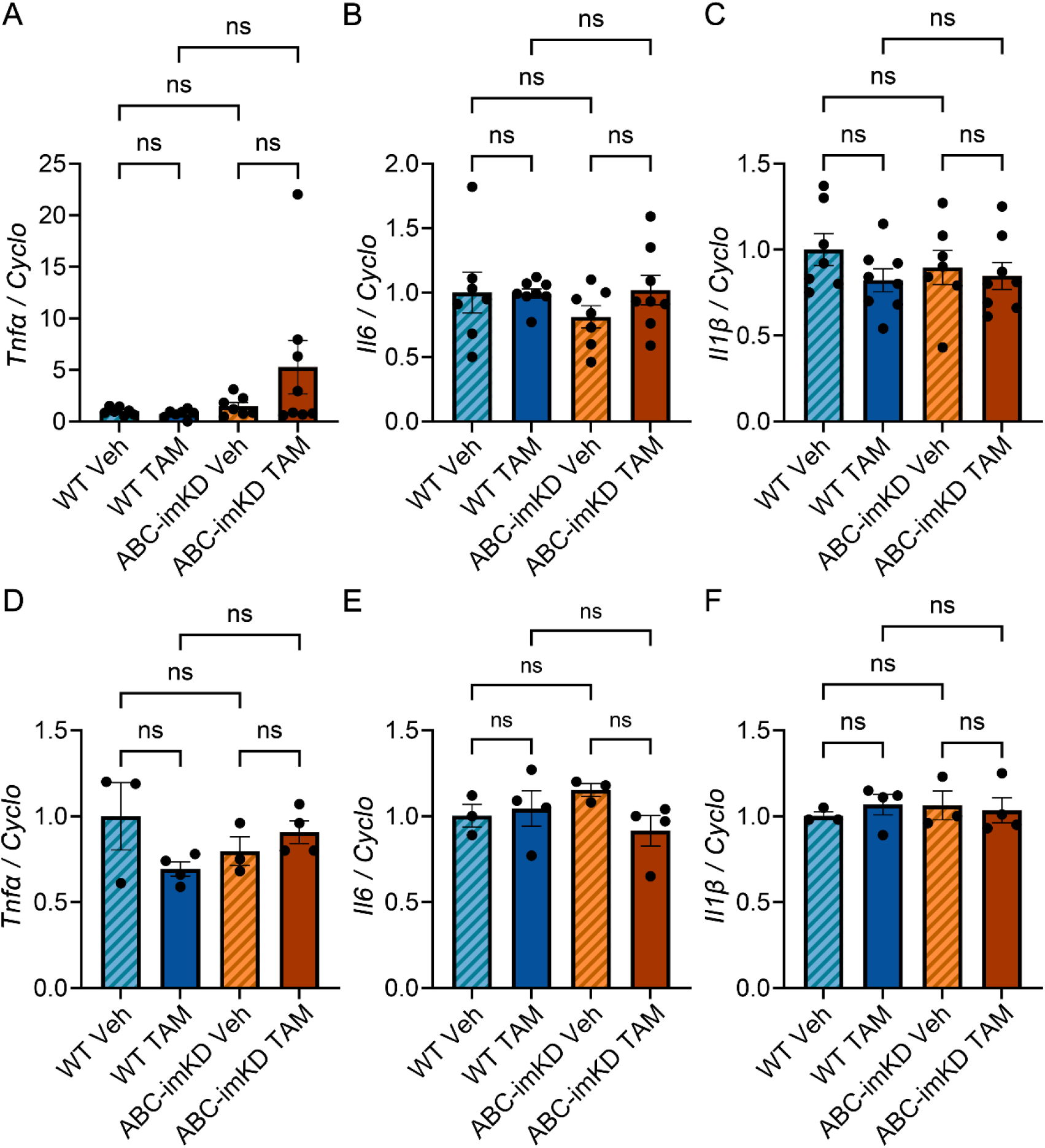
Expression of the pro-inflammatory cytokines *Tnfα*, *Il6*, or *Il1β* in the P12 hippocampus and cortex from WT and ABC-imKD mice. Relative *Tnfα* (**A**), *Il6* (**B**), and *Il1β* (**C**) mRNA expression (normalized to *Cyclo*) in the P12 hippocampus of vehicle- and tamoxifen-treated WT mice (Veh, *n* = 7; TAM, *n* = 8) and ABC-imKD mice (Veh, *n* = 7; TAM, *n* = 8). Relative *Tnfα* (**D**), *Il6* (**E**), and *Il1β* (**F**) mRNA expression (normalized to *Cyclo*) in the P12 cortex of vehicle- and tamoxifen-treated WT mice (Veh, *n* = 3; TAM, *n* = 4) and ABC-imKD mice (Veh, *n* = 3; TAM, *n* = 4). ns, not significant. Data are presented as mean ± SEM. SEM, standard error of the mean. Statistical analyses were performed using a Kruskal–Wallis test followed by Dunn’s multiple comparisons test (**A, D,** and **F**), or ordinary one-way ANOVA followed by Tukey’s multiple comparisons test (**B**, **C**, and **E**). *n*, number of mice.

